# Real-Time Brain-Machine Interface Achieves High-Velocity Prosthetic Finger Movements using a Biologically-Inspired Neural Network Decoder

**DOI:** 10.1101/2021.08.29.456981

**Authors:** Matthew S. Willsey, Samuel R. Nason, Scott R. Ensel, Hisham Temmar, Matthew J. Mender, Joseph T. Costello, Parag G. Patil, Cynthia A. Chestek

## Abstract

Despite the rapid progress and interest in brain-machine interfaces that restore motor function, the performance of prosthetic fingers and limbs has yet to mimic native function. The algorithm that converts brain signals to a control signal for the prosthetic device is one of the limitations in achieving rapid and realistic finger movements. To achieve more realistic finger movements, we developed a shallow feed-forward neural network, loosely inspired by the biological neural pathway, to decode real-time two-degree-of-freedom finger movements. Using a two-step training method, a recalibrated feedback intention–trained (ReFIT) neural network achieved a higher throughput with higher finger velocities and more natural appearing finger movements than the ReFIT Kalman filter, which represents the current standard. The neural network decoders introduced herein are the first to demonstrate real-time decoding of continuous movements at a level superior to the current state-of-the-art and could provide a starting point to using neural networks for the development of more naturalistic brain-controlled prostheses.

## Introduction

Brain-machine interfaces (BMIs) offer hope to the very high numbers of Americans (∼1.7%) with sensorimotor impairments^1^. To this end, cortical BMIs have allowed human patients using brain-controlled robot arms to perform a variety of motor tasks such as bringing a drink to the mouth^2^ or stacking cups^3^. Motor decoding algorithms are required to convert brain signals into a control signal, usually with position and velocity updates, for the prosthetic device. Despite the potentially non-linear relationship between neural activity and motor movements^4,5^, linear algorithms – including ridge regression, Kalman filtering, and Poisson processes – represent state-of-the-art performance in motor decoding^2,6-8^. Even with the rapid progress, many recognize that further developments are necessary to restore quick and naturalistic movements^2^.

Some gains in performance have already been achieved by adding non-linearities to classic linear decoders to leverage the likely non-linear relationship between neural activity and motor movements. For example, since neural activity is markedly different when moving compared to stationary postures, decoders have been introduced to move a prosthesis only when the desire to move is detected^4,7,9^. To leverage the non-linear relationship between kinematics and motor cortex neural activity, the classic Kalman filter has been adapted by expanding its state space^10^ or with Gaussian mixture models^11^ so that the algorithm can adopt different linear relationships in different movement contexts. In a particularly novel implementation, Sachs et al.^12^ implemented a weighted combination of two Wiener filters trained for either fast movements or near-zero velocities so that continuously decoded velocities largely draw upon the fast Wiener filter at the beginning of the trial and the slow-movement filter as the cursor approaches the target. However, for many of these approaches, performance is improved only for very specific tasks, and a general-purpose nonlinear approach is lacking.

Artificial neural network decoders, with their capability to model complex non-linear relationships, have long been thought to hold tremendous promise for brain-machine interfaces. They may ultimately also represent the most biomimetic motor decoder to transform motor cortex activity to realistic motor movements. However, early neural network decoders, prior to recent advancements in hardware, toolboxes, and training methods, were not found to improve performance over standard linear methods when decoding continuous motor movements^13,14^. Many advanced techniques employing recurrent neural networks and variational inference techniques show great promise for predicting prosthetic kinematics from brain signals (in offline testing). However, these techniques are often employed to perform classification^15^, as opposed to continuous motor decoding, and not used in real-time control of prosthetic devices (in online testing), likely because of the computational complexity^16,17^. Sussillo et al.^18^, however, did demonstrate real-time control of a computer cursor with a recurrent neural network in a non-human primate implanted with motor cortex arrays. However, this did not outperform a ReFIT Kalman filter in the same animals^6,18^. George et al.^19^ demonstrated control of hand and finger movements in human amputees with peripheral nerve interfaces using a convolutional neural network but again did not outperform a linear Kalman filter.

In this work, for the first time, we demonstrate a ReFIT neural network for decoding brain activity to control random and continuous two-degrees-of-freedom movements in real time using Utah arrays in rhesus macaques. The ReFIT neural network is compared with the ReFIT Kalman filter, which we use to represent the current state-of-the-art in linear decoders. The ReFIT Kalman Filter, introduced by Gilja et al.^6^, is a two-step training process that first computes the weights of a classic Kalman filter and then modifies the weights when the prosthesis direction is not toward the actual target. In this study, we find that the ReFIT neural network decoder substantially outperforms our previous implementation of the ReFIT Kalman filter^20-22^ with >60% increase in throughput by utilizing high-velocity movements without compromising the ability to stop. This enables the use of shallow artificial networks, which may resemble biological motor pathways, for motor decoding applications and may be the bridge toward high-velocity, naturalistic robotic prostheses.

## Results

Two adult male rhesus macaques were implanted with Utah arrays (Blackrock Microsystems, Salt Lake City, Utah) in the hand area of the primary motor cortex (M1), as shown in Fig. 1a. The macaques were trained to sit in a chair and perform a finger target task in which a hand manipulandum was used to control virtual fingers on a computer screen in front of the animal. During online BMI experiments, spike-band power (SBP) was used as the neural feature. SBP is the time-averaged power in the 300-1000-Hz frequency band that provides a high signal-to-noise ratio correlate of the dominant single-unit spiking rate, and usually outperforms threshold crossings as a feature^23^. A two-degree-of-freedom finger task was previously developed by Nason et al.^21^, where the monkeys used two individual finger groups to acquire simultaneous targets along an arc. Monkey N used his index (D2) finger individually and his middle-ring-small (D3-5) fingers as a group, and Monkey W used D2 and D3 as one group and D4 and D5 as the second group. However, unlike the previous task using center-out targets, targets herein were acquired randomly to increase task difficulty. After a 400-trial calibration task, a decoder was trained to predict velocity of both finger groups, as shown in Fig. 1b. We have recently demonstrated online real-time decoding of these 2 degrees of freedom using a ReFIT Kalman filter^21^, and primarily compare our novel algorithm to that approach.

**Fig. 1.**
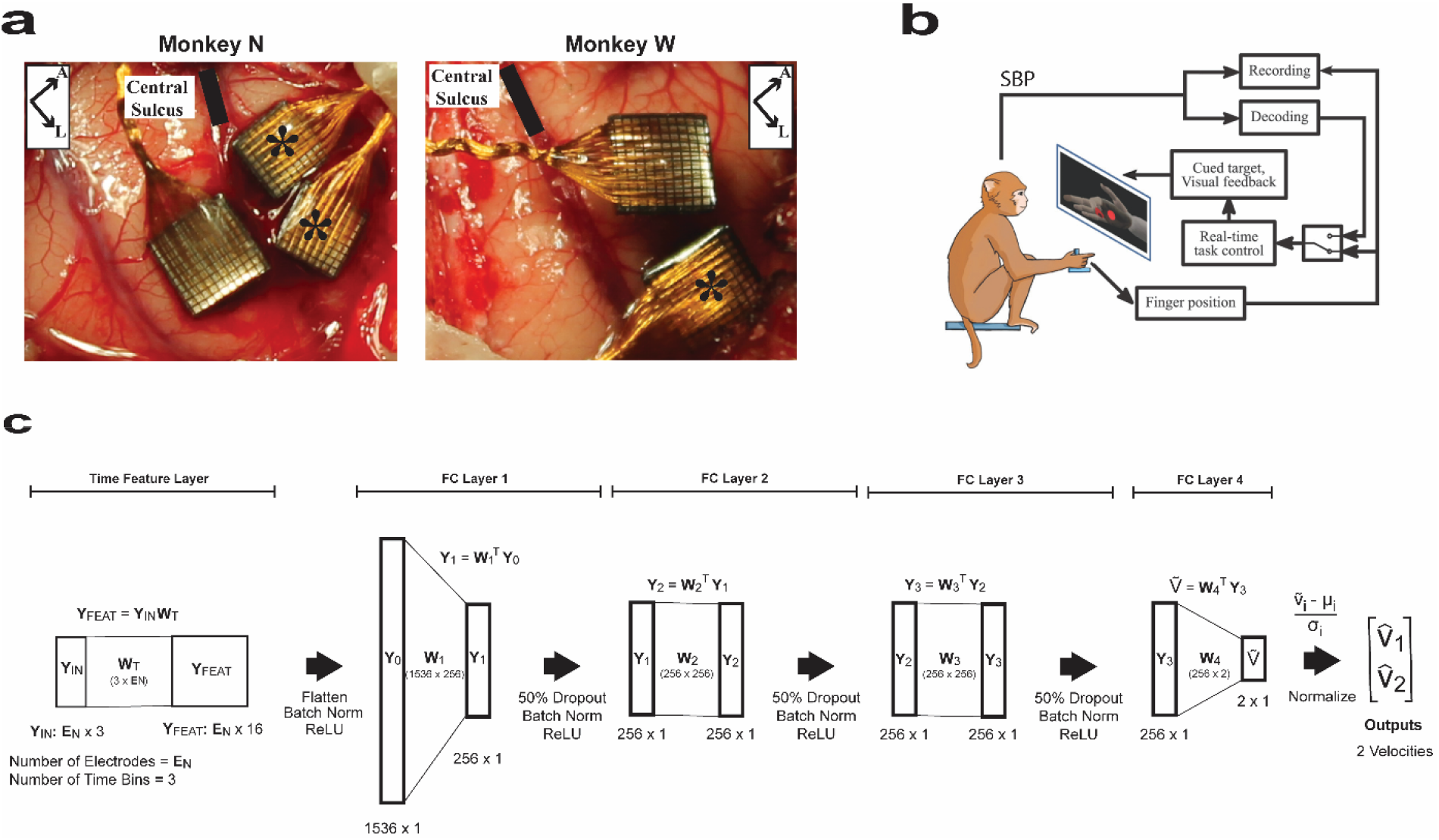
Neural network velocity decoder. **a**, Image of Utah array implants for Monkeys N (left) and W (right). In Monkey N, two split Utah arrays were implanted in primary motor cortex immediately anterior to the central sulcus and denoted with asterisks (*). The array in primary somatosensory cortex was not used in this analysis. In Monkey W, two 96-channel arrays were implanted and the analysis herein uses the lateral array. **b**, Experimental setup. The NHP is controlling the virtual finger with the hand manipulandum in manipulandum-control mode or using spike-band power (SBP) to control the virtual finger in brain-control mode. **c**, NN architecture. The network consists of five layers. The input to the network is ***Y***_***IN***_ that is a ***E***_***N***_ × 3 data matrix that corresponds to the number of input electrodes and the 3 previous 50-ms time bins. The time feature layer converts the last three 50-ms time bins for all the electrodes into 16 learned time features for each electrode. The equation representing the operation is given above the graphical description of the layer. The arrow indicates that the elements undergo batch normalization and pass through a ReLU function and are then flattened to a 1536 × 1 array. The remaining four layers are fully connected layers with an associated weight matrix, denoted by W. The first three layers consist of 256 hidden neurons and process the hidden neuron output first with 50% dropout, then batch normalization, and finally with a ReLU function. The fourth and final fully connected layer, FC-Layer 4, has two neurons – that are normalized – and represents final velocity estimates of the two fingers, 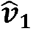 and 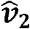 Panel **b** modified from Vaskov et al. and licensed under CC BY 4.0 (https://creativecommons.org/licenses/by/4.0/) / “spikes” replaced with SBP.

### Offline analysis of the neural network architecture

Limited computational complexity was a design goal for the neural network to allow same-day training and testing. As most online decoders incorporate recent time history^2,3^, the neural network was designed so that an initial time-feature layer constructed 16 time features per electrode from the preceding 150 ms of SBP (time feature layer in Fig. 1c). These time features were then input into 4 fully connected layers, where the first three output to a rectified linear unit (ReLU) activating function and the final layer outputs a velocity for each finger group. The number of fully connected layers and output time features were chosen to achieve near maximal correlation coefficient in offline performance using 400 trials of training data. As can be seen in Fig. 2a, increasing the number of neurons in hidden layers beyond 256 and the number of fully connected layers beyond 4 did not substantially improve the offline correlation. Furthermore, increasing the number of time features beyond 16 (Fig. 2b) also did not substantially improve offline correlation. For notational simplicity, the neural network in Fig. 1c is abbreviated as NN.

**Fig. 2.**
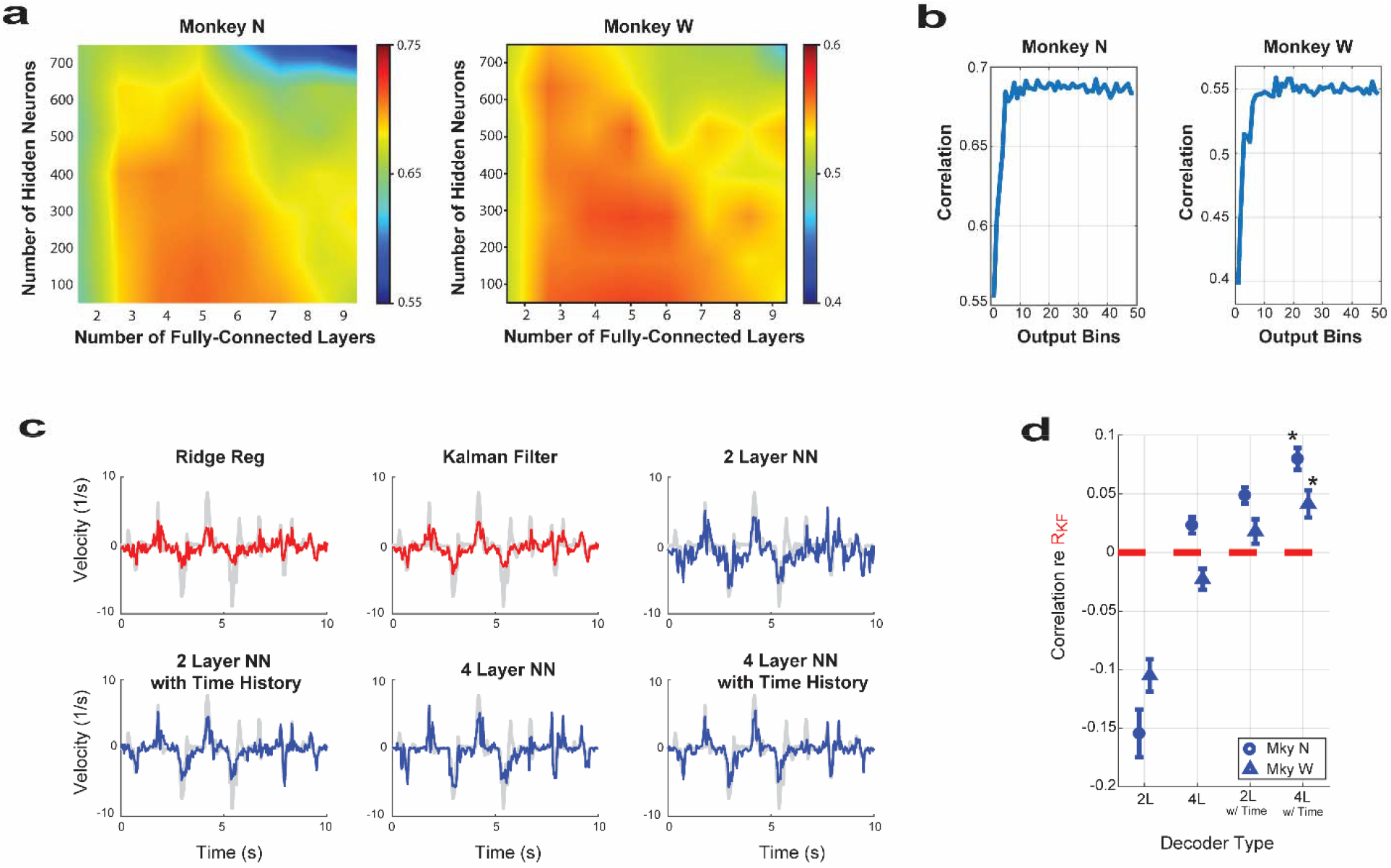
Neural network offline analyses. **a**, Heat map illustrating the offline correlation between the number of fully connected layers versus the number of hidden neurons for Monkeys N (left) and W (right). As in Fig 1c, the fully connected layers followed a preceding time feature layer. The correlation values were calculated as the average of the five-best correlation during the training of the network. The networks used during online testing had 4 layers and 256 neurons. **b**, The correlation during offline training for Monkeys N and W as a function of the number of learned time features in the output from the time-history layer of Fig. 1c. Four fully connected layers followed the time feature layer. The number of output features selected in online testing was 16. **c**, Examples comparing actual velocity (grey) and decoded velocity for linear decoders (red) and neural network decoders (blue) during 1 day of manipulandum-control tasks for Monkey N. The Y-axis is normalized by the standard deviation of actual velocities during the entire run. **d**, Mean (and S.E.M.) offline correlation difference between one of the neural network decoders and the Kalman filter, i.e., correlation of neural network decoder minus correlation of the Kalman filter. The circles denote the mean for Monkey N over both fingers over 3 d, and the triangles denote the mean for Monkey W over two fingers over 3 d. 2L = 2-layer neural network; 2L w/ Time = 2-layer neural network with a preceding time feature layer; 4L = 4-layer neural network; 4L w/ Time = 4-layer neural network with a preceding time feature layer. Asterisks (*) denote statistically significant differences between the correlation of the neural network decoder and the Kalman filter.

The impact on performance of the individual network components was assessed through an offline analysis based on 3 consecutive days of recorded spike-band power for each monkey during manipulandum-controlled finger task. Illustrative examples of predicted versus actual finger velocities for Monkey N using the manipulandum are given for neural networks of increasing complexity: 2 layers, 2 layers with time history, 4 layers, and 4 layers with time history (Fig. 2c). The correlation of each neural network decoder relative to the Kalman filter correlation is given in Fig. 2d by combining both fingers over all days for each monkey. The offline Kalman filter correlation averaged 0.59 ± 0.01 for Monkey N and 0.50 ± 0.02 for Monkey W. In both monkeys, the correlation is highest for the 4-layer network with time history (NN in Fig. 1c), followed by the 2-layer network with time history, followed by the 4-layer network without time history, followed by the 2-layer network without time history. In both monkeys, the 4-layer network with time history, NN, achieves a higher offline correlation than the Kalman filter (*P* = 3.6 × 10^−4^ for Monkey N and *P* = 0.016 for Monkey W), and the total performance comparisons are summarized in Table 1.

**Table 1:** Offline performance comparing Kalman filter and neural network (NN) decoders.

| Decoder | Correlation |  |
| --- | --- | --- |
|  | Monkey N | Monkey W |
| Kalman filter | $0.59 \pm 0.01$ | $0.50 \pm 0.02$ |
| 2-layer NN | $0.44 \pm 0.02$ | $0.40 \pm 0.01$ |
| 2-layer NN with time history | $0.64 \pm 0.01$ | $0.52 \pm 0.02$ |
| 4-layer NN | $0.61 \pm 0.01$ | $0.48 \pm 0.02$ |
| 4-layer NN with time history | $0.67 \pm 0.01$ | $0.54 \pm 0.02$ |

### Neural network decoder outperforms ReFIT Kalman filter decoder in real-time tests

In two non-human primates (NHP), Monkeys N and W, neural network decoders outperformed a ReFIT Kalman filter (RK) during real-time (online) testing, and the performance results are summarized in Table 2. In Monkey N, a neural network decoder outperformed the RK 13 mos after implantation in 2 days of testing over 1080 total trials, regardless of which algorithm was used first. The NN decoder improved the throughput over the RK by 26% with 2.15 ± 0.05 bits per second (bps) for the NN and 1.70 ± 0.03 bps for RK (*P* < 10^−5^). The acquisition time was 1240 ± 40 ms for the NN and 1550 ± 40 ms for the RK. NN had 3/543 unsuccessful trials while RK had 1/537 unsuccessful trial. In Monkey W, NN and RK decoders were compared 2 mos after implantation on one day testing over 412 trials. As graphically depicted in Fig. 3a, the NN decoder improved the throughput over the RK by 46%, with 1.23 ± 0.09 bps for the NN and 0.84 ± 0.04 bps for RK (*P* < 10^−5^). The acquisition time was 2680 ± 160 ms for NN and 3310 ± 130 ms for RK. NN had 26/133 unsuccessful trials while RK had 113/279 unsuccessful trials. Fig. 3a illustrates the throughput of each trial and the mean value for each run.

**Table 2:** Real-time performance comparison between ReFIT neural network (RN), neural network (NN), and ReFIT Kalman filter (RK) decoders. bps = bits per second.

|  | Monkey N |  |  | Monkey W |  |  |
| --- | --- | --- | --- | --- | --- | --- |
|  | RK | NN | RN | RK | NN | RN |
| **NN vs. RK** |  |  |  |  |  |  |
| Throughput (bps) | $1.70 \pm 0.04$ | $2.15 \pm 0.05$ | | $0.84 \pm 0.04$ | $1.23 \pm 0.09$ | |
| Acquisition time (ms) | $1550 \pm 40$ | $1240 \pm 40$ | | $3310 \pm 130$ | $2680 \pm 160$ | |
| Time to target (ms) | $1110 \pm 30$ | $950 \pm 20$ | | $2410 \pm 110$ | $1680 \pm 110$ | |
| Dwell time (ms) | $440 \pm 30$ | $290 \pm 20$ | | $790 \pm 100$ | $1000 \pm 120$ | |
| Successful (total) trials | 540 (543) | 536 (537) |  | 113 (279) | 107 (133) |  |
| **RN vs. NN** |  |  |  |  |  |  |
| Throughput (bps) | | $1.51 \pm 0.04$ | $2.15 \pm 0.05$ | | $1.20 \pm 0.06$ | $1.43 \pm 0.05$ |
| Acquisition time (ms) | | $1880 \pm 50$ | $1320 \pm 30$ | | $2610 \pm 120$ | $2220 \pm 70$ |
| Time to target (ms) | | $1230 \pm 30$ | $840 \pm 20$ | | $1740 \pm 80$ | $1430 \pm 50$ |
| Dwell time (ms) | | $650 \pm 30$ | $480 \pm 30$ | | $860 \pm 90$ | $780 \pm 60$ |
| Successful (total) trials |  | 616 (629) | 772 (772) |  | 185 (237) | 441 (483) |
| **RN vs. RK** |  |  |  |  |  |  |
| Throughput (bps) | $1.41 \pm 0.03$ | | $2.29 \pm 0.05$ | | | |
| Acquisition time (ms) | $1940 \pm 50$ | | $1270 \pm 30$ | | | |
| Time to target (ms) | $1330 \pm 30$ | | $790 \pm 10$ | | | |
| Dwell time (ms) | $610 \pm 40$ | | $470 \pm 30$ | | | |
| Successful (total) trials | 559 (614) |  | 737 (737) |  |  |  |

**Fig. 3.**
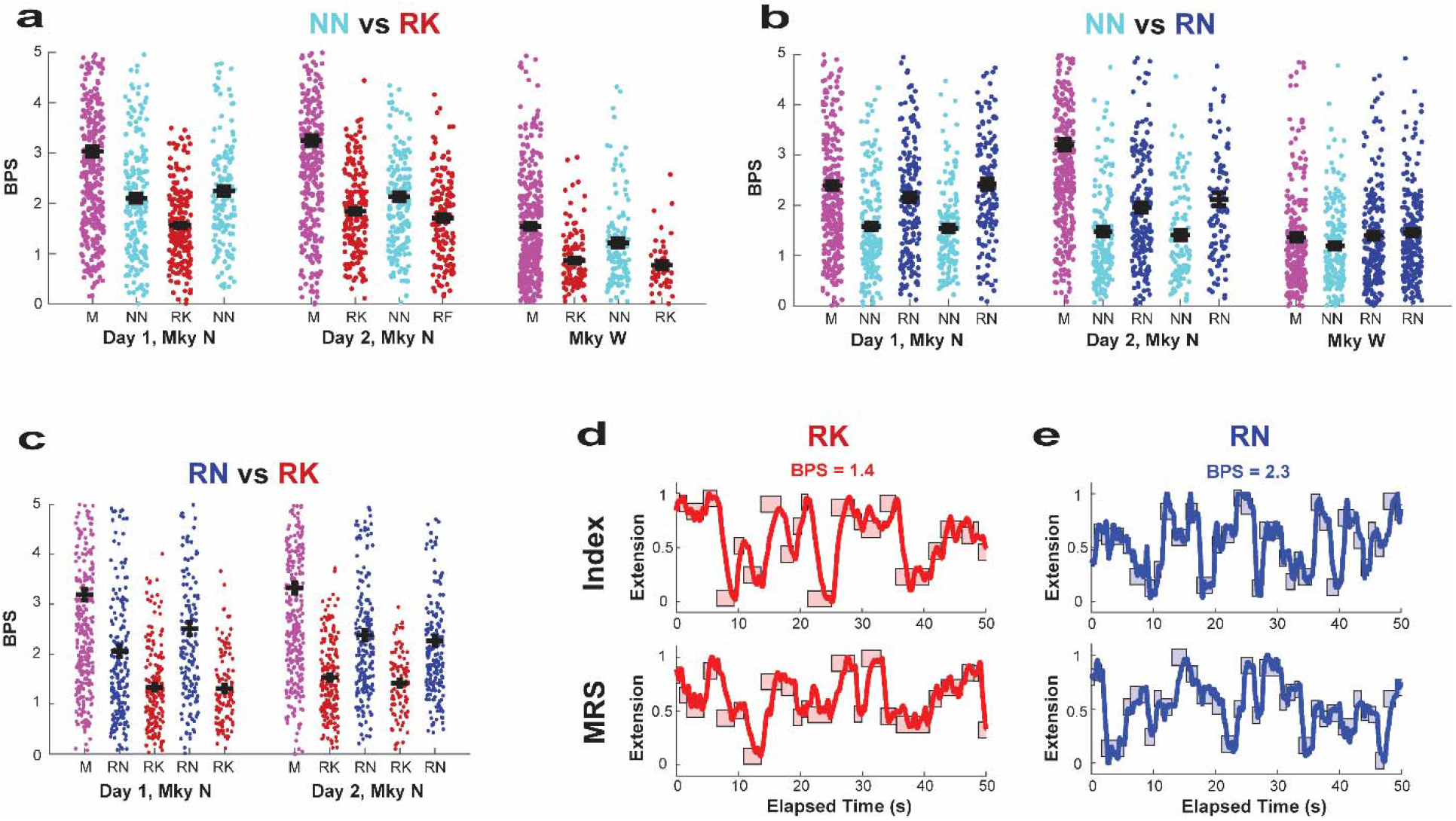
RN decoder outperforms RK during real-time tests. **a**, Throughput population data comparing the NN decoder (cyan) and the RK decoder (red) for 2 days of testing with Monkey N and 1 day of testing with Monkey W. The manipulandum-control runs are indicated with “M” (magenta). For each run, the throughput is indicated with a dot. The few values greater than 5 bps are not shown. The black bars represent the mean and S.E.M. **b**, Throughput population data comparing the NN decoder (cyan) and RN decoder (blue) for 2 days of testing with Monkey N and 1 day of testing with Monkey W. **c**, Throughput population data comparing the RN decoder (blue) and the RK decoder (red) for 2 days of testing with Monkey N. **d**,**e**, Raw decoded position using the RK (red; **d**) and RN (blue; **e**) for the index finger (top pane) and middle-ring-small (MRS) fingers in Monkey N, which are locked together (bottom pane). The targets are represented as the shaded box. The x-axis denotes the elapsed time, 50 sec, and the y-axis denotes the proportion of finger extension, i.e., 0 is fully extended and 1 is fully flexed. These time windows are representative of the average decoding performance as measured by throughput. M = manipulandum-control; NN = neural network; RK = ReFIT Kalman filter; RN = ReFIT neural network; BPS = bits per second.

### ReFIT neural network decoder outperforms both the original NN and RK decoders

The ReFIT innovation was applied to the neural network in a similar manner as it was used with the Kalman filter. Essentially, after completing trials using the NN decoder, the NN learned weights were further updated whenever the predicted finger direction was oriented away from the true targets, as described in the Methods. The ReFIT neural network (RN) decoder improved performance across all metrics when compared with the original NN in both monkeys (illustrated in Fig. 3b and Table 2). These tests for Monkey N were conducted at 19 mos post-implantation, and the decoding performance of all decoders had declined from earlier tests at 13 mos.

In Monkey N, who was capable of running a large number of consistent trials in 1 day, RN was compared directly with RK 19 mos after implantation in 2 days of testing with 1351 total trials (Fig. 3c and Table 2). RN improved the throughput over the RK by 62%, with 2.29 ± 0.05 bps for the RN and 1.41 ± 0.03 bps for RK (*P* < 10^−5^). Average performance of each decoder for the random finger task is illustrated in Videos 1 and 2. The acquisition time was 1270 ± 30 ms for the RN and 1940 ± 50 ms for the RK. There were no unsuccessful trials (0/737) using the RN and 55/614 unsuccessful trials with the RK. RN outperformed RK regardless of which algorithm was used first (*P* < 10^−5^), as illustrated in Fig. 3c in terms of throughput. Representative raw finger tracings are depicted in Fig. 3d for the RK and in Fig. 3e for the RN and depict a time segment with a throughput equivalent to the average throughput over both days. The tracings illustrate the higher target acquisition rate for the RN (30 targets in 50 sec) compared to the RK (21 targets in 50 sec).

### ReFIT neural network decoder outperforms optimized RK decoder

Our implementation of the ReFIT Kalman filter for finger control utilizes a physiological lag and does not include hyper-parameter tuning (i.e., gain and smoothing parameters)^20-22^. However, other work suggests RK performance can be improved without lag (providing the RK updates to the virtual fingers without delay)^24^ and by optimizing the online gain and smoothing parameters for RK^25^. In 2 days of testing at 29 mos post implantation with Monkey N, the zero-lag, optimized ReFIT KF, RK_opt_, was compared with RN in over 1164 trials using the same protocol as used above to compare RK and RN. In these tests, the throughput of RN of 2.41 ± 0.05 bps remained greater than that of RK_opt_ at 2.12 ± 0.05 bps (*P* = 1.2 × 10^−5^).

The performance of RK, prior to optimization, had also improved at 29 mos (around 1.91 bps on a separate day of testing). Thus, most (>50%) of the improvement of RK_opt_ over the 1.41 bps using RK at 19 months, as described in the section above, may be from improved NHP behavior with 10 additional months of practice with RK. For detailed evaluation of the performance improvement with RK_opt_ over RK, see “Optimizing the lag, gain, and scaling factors for real-time tests” in Materials and Methods.

### Neural network decoders allow higher velocity decodes than the Kalman filter

To better understand why the RN outperformed the RK decoder, the mean velocity over all the successful trials was computed for each decoder for both monkeys. As seen in Figs. 4a,b, virtual fingers controlled by the RN and NN decoders achieved higher peak velocities and were more responsive for both monkeys than when the virtual fingers were controlled by the RK decoder. For Monkey N (Fig. 4e), the time to the velocity peak averaged 300 ms for RN, 350 ms for NN, and 450 ms for RK. The peak of the averaged velocity was 1.35 ± 0.03 u/sec for RN, 1.00 ± 0.03 u/sec for NN, and 0.55 ± 0.02 u/sec for RK, where u denotes arbitrary units such that 1 was full flexion and 0 was full extension. For Monkey W (Fig. 4a), the time to peak was 350 ms for RN, 400 ms for NN, and 800 ms for RK. The peak average velocity was 0.94 ± 0.04 u/sec for RN, 0.76 ± 0.04 u/sec for NN, and 0.39 ± 0.04 for RK u/sec. Thus, in both monkeys, the average time to peak velocity and the peak velocity itself were improved with RN and NN than for the standard RK decoder. The high velocities achieved using RN is illustrated for a center-out task in Video 3.

**Fig. 4.**
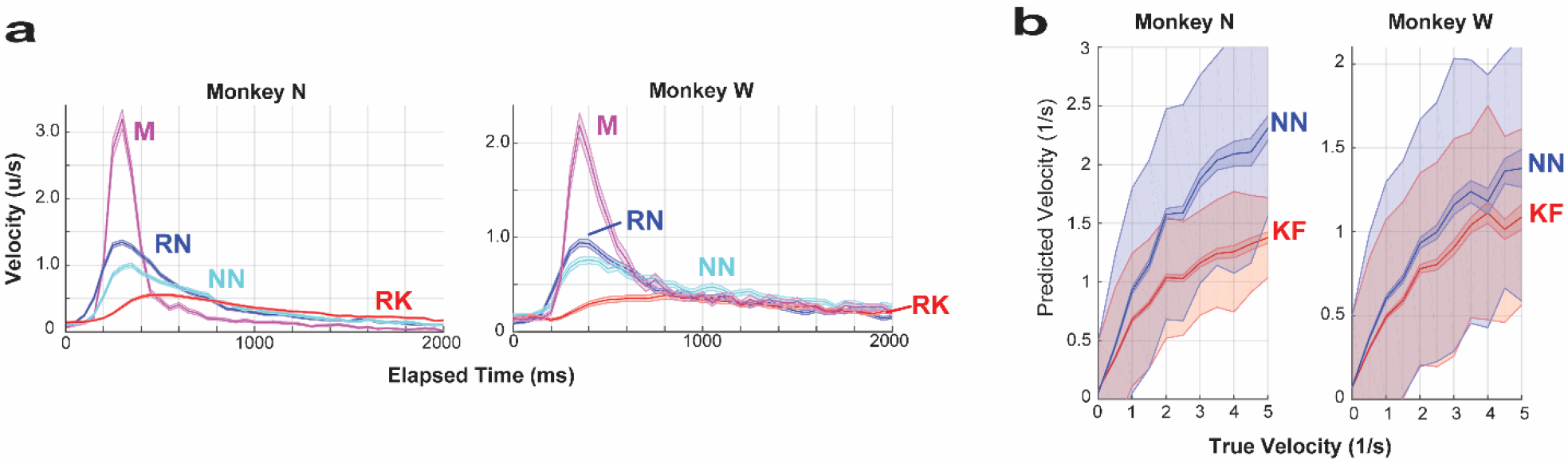
Higher decoded velocities using neural network decoders. **a**, Online analysis. Virtual finger velocity for Monkey N (left pane) and Monkey W (right pane) for RN (blue), NN (cyan), RK (red), and hand control (magenta). The plots indicate that the neural network decoders achieve higher peak velocities in real-time tests. In Monkey N, the RN, RK, and hand-control data are taken from the days comparing RN vs. RK, while NN velocities were derived from the day comparing NN and RK. In Monkey W, RN data were derived from the day comparing RN vs. NN. RK was derived from the day comparing NN vs. RK. Hand control and NN were derived from both days. The solid line indicates the mean value and the shaded region denotes the S.E.M. The shaded line tightly surrounds the mean, making these difficult to distinguish. If the trial was completed in less than 2000 ms, a velocity of zero was assigned extending from trial completion to 2000 ms. The unit, u, denotes arbitrary distance such that 1 was full flexion and 0 full extension. **b**, Offline analysis of true and predicted velocities for Monkey N (left) and Monkey W (right). The NN (blue) decodes higher velocities than the KF (red). The predicted velocity is normalized at a true velocity of zero so that one positive standard deviation of predicted velocities for both NN and KF is normalized to 0.5. The solid line indicates the mean value and the shaded region around denotes the S.E.M. The larger shaded region around the mean denotes the variance. NN = neural network; RK = ReFIT Kalman filter; RN = ReFIT neural network; KF = Kalman filter.

To ensure the NN does not achieve high-velocity decodes at the expense of low-velocity decoding accuracy, which is important for stopping the prosthesis, the predicted velocity as a function of the true velocity was compared for the NN and Kalman filter (KF) in an offline analysis (graphically shown in Fig. 4b). The predicted velocity on the vertical axis is scaled so that when the true velocity is at zero, the standard deviation of the predicted velocity equals 1/2. In the high-velocity range (>1 u/s), the decoded NN velocity averages 157 ± 3% of the KF velocity for Monkey N and 122 ± 2% for Monkey W. Thus, after accounting for decoder performance at low velocities, the range of velocities that can be achieved is higher for the NN than the KF. As shown in the online analysis, higher velocities improved the performance of NN decoders.

### Neural network merges decoders optimized for positive and negative velocities

Due to the network architecture itself, each node of the final hidden layer contributes either a positive or a negative velocity to the final prosthetic finger velocity. We explored whether this itself provides an example of how the fit is improved for different movement contexts, i.e. positive and negative velocities. Specifically, for finger 1, the sum of the product of ***N***_***k***_ and ***W***_***4***_^***(1***,***k)***^, over all *k*, determine 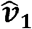, where ***N***_***k***_ is the *k*-th node of the final hidden layer and ***W***_***4***_^***(1***,***k)***^, represents the learned weights (shown in Fig. 5a). Since each ***N***_***k***_ is the output of the ReLU function, ***N***_***k***_ is necessarily greater than or equal to zero. Thus, the *nodal contribution* of the k^th^ node, 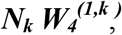 can be either positive (if ***W***_***4***_^***(1***,***k)***^ > 0) or negative (***W***_***4***_^***(1***,***k)***^ < 0) – but not both positive and negative.

**Fig. 5.**
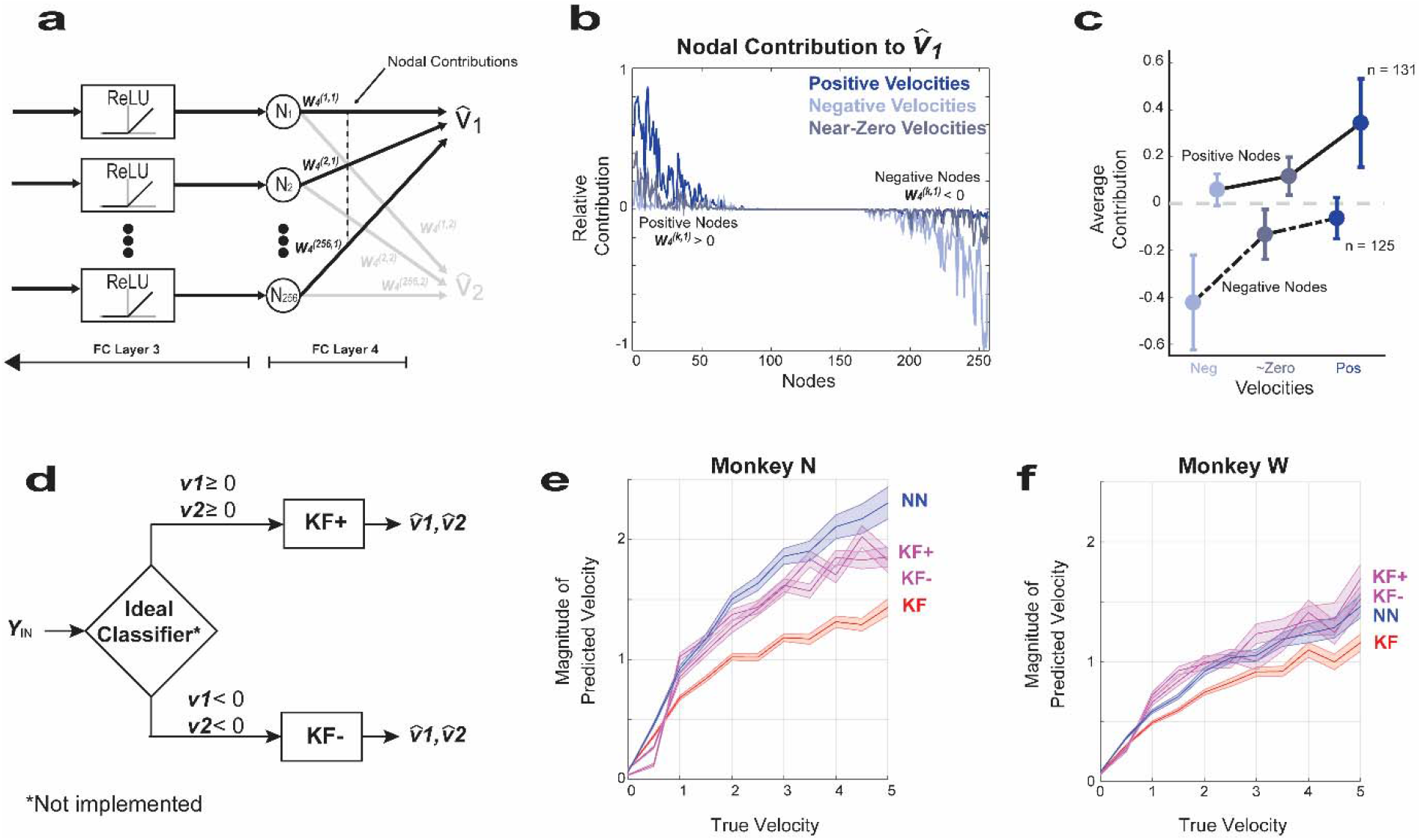
Neural network decoder functions as a weighted combination of a positive and negative velocity decoder. **a**, Final neural network layer that converts 256 nodes into 2 finger velocities by matrix multiplication by a matrix ***W***_***4***_^***(i***,***j)***^ of size 256 × 2, where ***i*** denotes the row and ***j*** denotes the column value. The value of the 256 nodes, ***N***_***k***_, of the final layer are all positive given that they are derived from the output of the preceding ReLU function. For finger 1, the ***k***^th^ node was considered a “positive node” if it contributes to positive – not negative – velocities and occurs when ***W***_***4***_^***(K***,***1)***^ > 0. “Negative nodes” contribute negative – not positive – velocities when ***W***_***4***_^***(K***,***1)***^ < 0. The nodal contribution from the ***k***^th^ node is defined as ***N***_***k***_***W***_***4***_^***(K***,***1)***^ and is the value of the output at the dashed line. **b**, Example illustration of the mean nodal contribution from positive and negative nodes when the true velocity is positive (dark blue), negative (light blue), and near zero (grey) during 1 day of testing with Monkey N. As illustrated in the figure, positive nodes are much higher than negative nodes during positive velocities, while negative nodes have a higher magnitude than positive nodes during negative velocities. Positive velocity is defined as ***v***_***1***_ > σ, negative velocity is defined as ***v***_***1***_ < -σ, and velocities near zero are defined as -σ/4 < ***v***_***1***_ < σ/4, where σ is the standard deviation of true finger velocity. **c**, The nodal contribution during positive, negative, and near-zero true velocities illustrates that positive nodes largely determine the numerical value of positive velocities and negative nodes largely determine the value during negative velocities. The nodal contribution is averaged across both fingers during the 3 offline days for both Monkeys N and W. Thus, three operating regimes exist for the neural network decoder that consist of a decoder during positive velocities based on positive nodes, a decoder during negative velocities based on negative nodes, and a decoder using positive and negative nodes during velocities near zero. The dots indicate the mean value and the error bars indicate the standard deviation. **d**, Block diagram of a hypothetical decoder that uses two separate filters for positive finger velocities, ***v***_***1***_ ≥ 0 and ***v***_***2***_ ≥ 0, and negative finger velocities, ***v***_***1***_ < 0 and ***v***_***2***_ < 0. The Kalman filter for positive velocities, KF(+), was trained on velocities near zero and positive values (> -0.5σ), whereas the Kalman filter for negative velocities, KF(-), was trained on mainly negative velocities (< 0.5σ). The ideal classifier is depicted only to illustrate the concept and was not implemented. **e**,**f**, True versus predicted velocity magnitude during 3 days of manipulandum-control testing (3 days each for Monkeys N and W) for the neural network decoder (blue), Kalman filter (red), KF(+) (magenta), and KF(-) (magenta). The full-range Kalman filter predicted both positive and negative velocities, KF(+) predicted only positive velocities, and KF(-) predicted only negative velocities. The magnitude estimated velocities of KF(+) and KF(-) are shown to be higher than those of the full-range Kalman filter and illustrate that training and implementing the Kalman filter over restricted ranges would allow for higher velocities (assuming an ideal classifier). The solid line indicates the mean value and shaded lines indicated the S.E.M.

The nodal contributions of positive and negative nodes during day 1 for Monkey N are illustrated in Fig. 5b for positive velocities (***v***_***1***_ > σ), negative velocities (***v***_***1***_ < σ), and near-zero velocities (-σ/4 < ***v***_***1***_ < σ/4). During positive velocities, the final estimate, 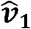, of ***v***_***1***_ is dominated by positive nodes, which is illustrated with the dark blue line that depicts much higher nodal contributions for positive than negative nodes. The same is true of negative nodes during negative velocities. This trend is confirmed at a population level for both monkeys in Fig. 5c, where positive nodes dominate negative nodes at positive velocities and vice versa for negative velocities. Thus, the NN decoder is capable of optimizing for positive velocities by learning weights of positive nodes and optimizing for negative velocities through the weights of negative nodes.

To understand whether the network is improving on the Kalman filter via separating movement contexts, we trained two separate Kalman filters: one for positive velocities (***KF+***) and one for negative velocities (***KF-***), as illustrated in Fig. 5d. The decoder in Fig. 5d assumes a perfect classifier that correctly chooses either ***KF+*** or ***KF-***depending on whether the true velocities are positive or negative. As can be seen for both monkeys in Figs. 5e,f, ***KF+*** and ***KF-***achieve higher velocity magnitudes closer to the NN decoder, and unlike the original Kalman filter, covers a wider range of velocities. This suggests that the NN allows for optimal fits within both of these contexts without overt switching.

## Discussion

During two-degree-of-freedom finger decoding, the ReFIT NN (RN) improved performance by more than 60% over our implementation of the ReFIT Kalman filter (RK) for finger control^20-22^. Even with RK parameter optimization and months of practice on using RK to control finger movements, RN continued to demonstrate a substantial performance advantage. This improvement was driven by more accurately decoding higher velocities. Accurate high-velocity decodes may arise by separately training weights for either positive or negative velocities, and this may lead to more robust performance in various real-life tasks.

### Performance improvement with neural network decoders

Neural network algorithms are loosely inspired by biological neural networks, and many have explored their use in brain-machine-interface decoding applications^13,14^. Although many sophisticated network architectures show tremendous promise in simulation^16,17^, neural network decoders have not improved real-time performance over standard linear techniques^13,14,18,19^. Our approach differs from previous approaches mainly by using a shallow network architecture and by leveraging the multi-step training procedure to improve training data originally developed for the ReFIT Kalman filter^26^. Additionally, we utilize recently developed regularization techniques to prevent overfitting (i.e., batch normalization and dropout)^27-29^ and also incorporate 150-ms time history of spiking band power (SBP) as input to the decoder instead of only one point in time.

The shallow network architecture of 1 time-feature layer and 4 fully connected layers allows for the roughly 5 × 10^5^ learned parameters to be trained with only 400 trials of same-day training data in about 1 minute. In contrast, recurrent neural network architectures have combined training data across multiple days^15,16^, and when implemented as continuous motor decoders may be too complex to run in real time. An additional benefit of shallow feed-forward networks is that computing the velocity in real-time mode introduces only a 1-2 ms lag. Thus, through use of a shallow network, limitations typical of more computationally expensive architectures are avoided.

Incorporating a two-step, intention-based, re-training step is the fundamental innovation improving the ReFIT over the classic velocity Kalman filter^6^ and appears to have a similar positive effect on the NN. The intention-based retraining step was already known to improve finger decoding for a Kalman filter during center-out tasks^22^. Similar to the substantial improvement seen in the ReFIT Kalman filter^6^, retraining the ReFIT neural network resulted in a substantial 50% improvement in performance over the original neural network decoder.

### Neural network decoder learns positive and negative velocity features for finger movement

By training the neural network, the learned weights for either positive or negative nodes appear to be optimized for either the positive or negative velocity range. Similar to our results, Sachs et al.^12^ showed that splitting the full velocity range into intervals subserved by separate Wiener filter decoders fine-tuned for either high or low velocities improved brain-machine interfaces for cursor control. Additionally, Kao et al.^7^ improved performance by 4.2-13.9% over the ReFIT Kalman filter using a hidden Markov model to enable movement only when the decoder is in a “movement state,” as neural activity is known to be different in movement and postural states^4^. The neural network architecture may be better able to discover these contexts without explicit classifiers or supervised training.

The variation of decoded velocities of the neural network appears to more closely mimic the range of velocities seen in native finger movements. In a tantalizing hypothesis, the decoder’s naturalistic movements may be related to its shallow architecture, which may resemble true biological pathways. Specifically, there are only a few synapses between the neurons in motor cortex and the α-motor neurons in the anterior horn of the spinal cord^30^. Although speculative, neural network architectures may perform well partly because motor cortex activity naturally controls flexion and extension of antagonist muscle pairs. The neural network architecture may readily decode this flexion and extension into positive and negative velocities. Whether the similarity of neural network decoders to biologic network leads to more naturalistic motor control of many more contexts and simultaneous degrees of freedom awaits further study.

## Limitations

As opposed to using our typical center-out task finger task^21^, we increased the difficulty to better challenge the decoders and elucidate differences between two well-performing decoders. Although these results could apply to a variety of real-word finger tasks, other tasks and prostheses (i.e., robotic arms) were not explicitly tested. While improvements in the Kalman filter or its implementation may increase its performance, the neural network performance can also be further optimized, such as by including position information into the decoder, optimally tuning its parameters, or by implementing it in a form similar to the steady-state Kalman filter by merging historical velocity and current updates (Eq. 2). Regardless, in online tests, the RN was found across all metrics to outperform RK, and offline tests confirmed a superior dimensionality reduction (as measured by correlation coefficient) and a higher dynamic range of predicted velocities. A piecewise implementation of the Kalman filter could conceivably be used to achieve a similar range of predicted velocities but would require a sophisticated and generalizable switching algorithm to choose the appropriate Kalman filter. Furthermore, the neural network algorithms could similarly be constructed for distinct velocity ranges. Lastly, while the neural network does require increased computational complexity, it was optimized for performance and not to optimally trade off performance with computational complexity, which could certainly be accomplished.

## Conclusion

This novel neural network decoder outperforms a current state-of-the-art motor decoder and achieves movements similar to naturalistic finger control. The architecture in large part resembles biological motor pathways and may be amenable to further performance improvements.

## Materials and Methods

### Implantation procedure

The protocols herein were approved by the Institutional Animal Care and Use Committee at the University of Michigan. Two adult male rhesus macaques were implanted with Utah arrays (Blackrock Microsystems, Salt Lake City, Utah) in the primary motor cortex (M1). Under general anesthesia and sterile conditions, a craniotomy was made and M1 was exposed using standard neurosurgical techniques. The arcuate sulcus of M1 was visually identified, and the array was placed where this sulcus touches motor cortex (Fig. 1a), which we have previously used as a landmark of hand area in rhesus macaques. The incision was closed, and routine post-anesthesia care was administered.

### Experimental setup and finger task

Both Monkeys N and W were trained to sit in a monkey chair (Crist Instrument, http://www.cristinstrument.com), with their head secured in customized titanium posts (Crist Instrument), while the Utah array was connected to the Cerebus neural signal processor (NSP, Blackrock Microsystems). The arms were secured in acrylic restraints. The hand contralateral to the motor cortex implant was placed in a manipulandum, described by Vaskov et al.^22^, that translates finger position to a number between 0 (full extension) and 1 (full flexion). A computer monitor was in plain sight for the NHP and depicted a large virtual hand (Fig. 1b). The virtual finger could be controlled in either manipulandum-control mode or in brain-control mode (i.e., brain signals converted to updates for virtual finger). Brain-control mode is commonly denoted as either real-time, closed-loop, or “online” mode. Manipulandum-control mode is often described as “offline” mode. The two-dimensional finger task is identical to the task developed by Nason et al., except performed on random instead of center-out targets^21^. The finger task required placing either the virtual index and/or ring finger on the target for 750 ms during training mode and 500 ms during testing mode (testing vs. training modes will be explained in a subsequent section). The target size was 15% of the active range of motion. With target acquisition, apple juice was automatically administered through a tube placed in the animal’s mouth.

### Front-end processing

The Utah array was connected to the Cerebus NSP (Blackrock Microsystems) through a cable. Although 96 channels were available, only channels that were not artifactual and with morphological neural spikes on the day of experiments or had shown morphological spikes in the past were included, leaving 54-64 channels for Monkey N and 50-53 channels for Monkey W. The Cerebus system sampled data at 30 kHz, filtered it to 300-1000 Hz, down-sampled it to 2 kHz, then transmitted it to the xPC Target environment (Mathworks, Natick, MA). The xPC Target computer took the absolute value of the incoming data and then calculated each channel’s mean in regular 50-ms time intervals. This binned value is referred to as spike-band power. We have previously shown that this band is highly correlated with and specific to the spiking rate of single units near the recording electrode^23^.

### Software architecture

A separate computer with one 2070 super NVIDIA GPUs (NVIDIA, Santa Clara, CA) was connected to the xPC. This computing box was called the e**X**ternal **G**raphic **P**rocessing **C**omputer (xGPC). The xGPC executed commands in Python (v3.7, https://www.python.org/) using the PyTorch library (v1.4, https://pytorch.org/). Real-time performance was guaranteed in the following fashion. The xPC transmitted data to the xGPC with a timestamp, the xGPC calculated updates for the virtual fingers from the inputs (for all decoders) and transmitted the data back to the xPC along with the original timestamp. When the xPC received the data packet, the packet was logged with a new timestamp. Real-time performance was guaranteed given that the timestamp received from xGPC (the original timestamp sent by xPC) was within 50 ms of the current xPC timestamp and updates to the virtual fingers occurred every 50-ms time bin.

### ReFIT Kalman filter

The ReFIT Kalman filter (RK) was implemented for use with fingers as described by Vaskov et al. and Nason et al.^21,22^ In summary, it is a two-step process that involves first training a Kalman filter (KF) using spike-band power measurements from any 96 channels of the Utah array to predict updates to position and velocity states of the virtual fingers. A detailed description on the KF implementation is described by Vaskov et al^22^.

The trained KF was then used to perform closed-loop motor decoding. To train the RK, the target position for each finger is mapped to a two-dimensional space and the true velocity of each finger is scaled to be proportional to each finger’s distance to the target while keeping the total velocity magnitude constant. This method of ReFIT was introduced by Nason et al.^21^ and was not found to be statistically different from the ReFIT method in Vaskov et al.^22^, where the finger velocity was modified by multiplying velocities by -1 when the velocity was oriented in the opposite direction as the target. The KF was then retrained using these new velocity values (for details see Nason et al.^21^). As also detailed in Vaskov et al.^22^, Kalman gain was implemented with no position uncertainty.

Optimal lag is commonly implemented in KF motor decoders^31^ to account for the physiologic lag between cortical activity and motor movement^32^. Thus an optimal time lag, calculated to be one 50-ms bin for both Monkeys N and W, was applied when training and implementing the KF, as detailed in previous work^20,22^. Control tests comparing zero and one 50-ms bin lag are provided below (see Section “Optimizing the lag, gain, and scaling factors for real-time tests”). Additionally, the Kalman filter can be implemented as a steady-state Kalman filter with a gain and smoothing factor that can be optimized for online tests. Our implementation generally does not tune these parameters as the ReFIT training algorithm may determine near optimal values for these parameters^33^. To validate this simplification, we compare our implementation of RK with one with optimal tuning (see Section “Optimizing the lag, gain, and scaling factors for real-time tests”). As will be explained below, we did compare RN with RK_opt_, which uses zero lag and optimally determined gain/smoothing parameters, to ensure our results hold against a theoretically optimized RK, with the results presented in the section “ReFIT neural network decoder outperforms optimized RK decoder” of Results.

To determine whether the Kalman filter could better predict the high velocities if trained and used on restricted velocity ranges, we conducted an offline analysis using “KF+” and “KF-.” KF+ was calculated with only positive and near-zero velocities, i.e., velocities greater than -σ/2, and KF-was calculated with velocities less than σ/2. These Kalman filters were trained as described by Wu et al.^31^ with position uncertainty and using the optimal physiologic lag calculated on that day.

### Neural network velocity decoder

The neural network velocity decoder was designed from preliminary offline experiments that explored various network architectures. The final network is given in Fig. 1c. The first layer was the time feature layer that constructs time features from 150 ms (three 50-ms bins) from the input electrodes. This layer was implemented in Pytorch, using the *torch*.*nn*.*Conv1d* module, i.e., as a one-dimensional convolution with a kernel size of 1 (H=W=1) and 3 input channels (neural network channels, not electrode channels). Each channel corresponded to one 50-ms time bin. Although possible to construct a spatial convolution across electrodes, this was not performed because the spacing between electrodes was hypothesized to be distant relative to the size of the neurons being recorded. The output of the time feature layer provided 16 features per electrode and, when flattened, provided 1536 inputs to a series of fully connected layers. Regularization for fully connected layers 1-3 included 50% dropout^27^ and batch normalization^28^. Fully connected layer 1 converted the 1536 inputs to 256, and the remaining layers had 256 hidden neurons. The sequence of the modules used was *torch*.*nn*.*linear, torch*.*nn*.*Dropout, torch*.*nn*.*BatchNorm1d*, and then finally *torch*.*nn*.*functional*.*relu*. The final layer implemented a matrix multiplication with *torch*.*nn*.*linear* to convert the 256 inputs to the two velocity estimates. The output of the network was normalized to zero mean and unit variance and roughly twenty times the magnitude of actual velocity peaks. This normalization was discovered to converge more quickly when training the RN than training without the normalization. The output of the neural network was scaled by an unlearned gain factor that equaled the average magnitude peaks of the actual velocity divided by the average magnitude peaks of the predicted velocity. No offset was applied to the final predicted velocity, leaving it a zero-mean signal. A diagram of the final neural network is given in Fig. 1c.

Prior to training the neural network, a training data set was collected in manipulandum-control mode for roughly 400 trials with randomly appearing targets. A subsequent 100 trials were also performed and served as a validation set to ensure the network had converged. This validation set was also used to calculate the gain as described above. If there was a non-zero median, this was subtracted as well to approximate a zero-mean signal. The SBP and velocity data were assembled into data structures in Matlab (Mathworks). The data were randomized in two ways. First, the time data were randomized into batches of 64 × 3 time points: 64 time points with the corresponding value at time delays of 0, 50, and 100 ms. Second, a triangular distribution of velocities was imposed on the training data spanning the range of -4σ to 4σ, where σ was the standard deviation of the actual velocity. A total of 20,000 training samples were randomly chosen to achieve this velocity distribution. In preliminary experiments, this velocity redistribution was observed to improve performance on the finger task when the neural network was trained on a center-out finger task that led to a “sticky finger” behavior, in which the finger would often get close to but not quite all the way to the target. However, when the neural network was trained on random targets, the velocity redistribution was not observed to improve performance over non-redistributed data, but we describe it here for completeness. This redistribution of velocities was also used when training on random finger targets so that the decoder could easily be generalized to other training paradigms in the future.

In addition to the neural network used for online testing, several other neural networks were used to understand how individual components of the neural network affected offline performance. The networks included a network of only two layers and no regularization (no batch normalization, dropout, or output normalization). There were also regularized networks, including a 2-layer fully connected network (256 hidden neurons) with a preceding time feature layer (3 input channels for each electrode and 16 output channels), and 4-layer fully connected network (256 neurons) with a time feature layer. These networks included regularization and parameters similar to Fig. 1c. The offline networks were compared with the classic Kalman filter (without the intention retraining step) and ridge linear regression without time history and with a regularization constant of λ = 10^−4^.

When training the network for online decoding, the neural network was optimized over 3500 iterations using the Adam optimization algorithm^34^ with a learning rate of 10^−4^, weight decay of 10^−2^, and momentum of 0.9. Each iteration consisted of a 64×3 mini batch (64 random time steps with 3 samples of 150 ms of time history). We attempted to use a relatively large learning rate as larger learning rates provide additional regularization for the network^29^. On one day for Monkey W, 3000 iterations were used. The number of iterations were determined for each network from the first of three offline testing days and chosen so that the correlation between actual and estimated velocity (on the testing set) did not significantly change with additional training iterations (changes in correlation with additional iterations on the order of ∼0.01). When generating weights for offline analysis, a learning weight of 2 × 10^−5^ allowed better comparisons between networks with different numbers of layers. Kaiming initialization was used to initialize the weights of each layer^35^, and the bias terms were initialized to zero. The dropout level used was 50%^27^. On each day, a training set (∼400 trials) and testing set (∼100 trials) were collected, and performance on a testing set was characterized by the correlation of predicted and actual velocity.

When searching for the preferred number of layers, hidden neurons, and output time features (Figs. 2a,b), performance was characterized by the average of the maximum of 5 correlations with the testing set over all training iterations. In this way, the optimal number of training iterations did not need to be calculated for different size networks.

The weights for the ReFIT neural network were calculated by first using the NN decoder in brain-control mode. A truth signal was then constructed from the original NN output by flipping the velocity direction whenever the estimated finger velocity was directed away from the target. Using this truth signal and the original neural network output, 500 further iterations of stochastic gradient descent were applied to further optimize the weights of the neural network. The gain factor for the neural network was calculated in the same manner as the original neural network except when comparing the NN to RN over 2 d, where the gain factor used during brain control with the NN was simply scaled by a factor 0.75 on both days.

### Testing protocols

Targets for the fingers were not allowed to be separated by greater than 50% of the range. During training, the random targets spanned 100% of the finger flexion/extension range, but during online decoding only 95% of the range was used. The classic Kalman filter was then used on a ∼250 trial run from which the ReFIT KF coefficients were calculated.

For Monkey N, 8 online testing days were conducted. Offline testing was performed using manipulandum-control trials from 3 consecutive days. Two of these days were the manipulandum-control training trials from the online testing comparing NN and RK conducted 13 mos post-implantation. An additional day of manipulandum-control data at 13 mos post-implantation was also included. For Monkey W, 2 online testing days were conducted. The offline analysis included manipulandum-control trials from 3 days at 2 mos post-implantation that included the 2 online days and an additional day of manipulandum-control trials. To reduce confounders when comparing decoders for each monkey, decoders were compared in an alternating lineup: either A-B-A or A-B-A-B testing. In 1 day for Monkey W, NN was compared with RN without alternating the decoder. For Monkey N, the first 50 trials with each decoder were discarded in the analysis. For Monkey W, only the first trial was discarded as there were fewer total trials since W was less motivated to complete trials. To visually illustrate the peak performance of the RN on our typical center-out task^21^, one additional day was included using the RN on this task.

### Performance assessment and statistical analysis

Performance in online mode was characterized with Fitt’s law throughput given below in Eq. 1, which accounts for both task difficulty and the time needed for completion. The variable *D*_*k*_ is the distance of the *k*-th virtual finger to the center of the *k*-th target at the start of the task, *S* is the target radius (equal in both fingers), and *t*_*acq*_ is the time to reach the target.

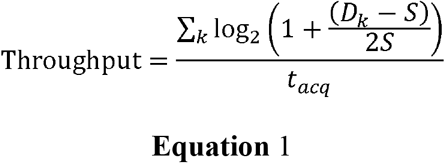

While the throughput was the primary performance metric, acquisition times were also reported.

All velocities in the offline analyses were normalized by the standard deviation of the true velocity with 1 indicating the equivalent of 1 standard deviation of the actual velocity. The time plots depicting the actual versus predicted velocity were selected from one of the training days to illustrate the results (Fig. 2a). The correlation for each decoder was averaged over 2 fingers on 3 days (Fig. 2b). The plots of true versus predicted velocity were calculated by binning the magnitude of the actual velocity into bins of size 1.0 at intervals of 0.5 and averaging the magnitude of the predicted velocity in each respective bin.

Comparing the performance metrics between groups was made with a paired t-test.

### Optimizing the lag, gain, and scaling factors for real-time tests

As explained above, we utilized physiologic lag similar to previous studies^31^ in our implementation of RK for finger control^20-22^. Unless otherwise mentioned, our comparisons of RK and RN use RK with a 50-ms bin lag. To evaluate the effect of a 50-ms bin lag, RK with a lag of one 50-ms bin was compared to RK with a zero-lag implementation (where the Kalman filter predictions update the virtual hand as soon as they are available) in one day of testing over 273 trials with Monkey N. The zero-lag RK improved throughput by 16% and peak average velocity by 13%.

The Kalman filter estimates of position and velocity can be simplified (assuming a steady-state Kalman gain) as a weighted sum of two components: the previous time step’s estimate of position/velocity and the current time step’s estimate of position/velocity derived from intra-cortical array as shown below in Eq. 2^25^.

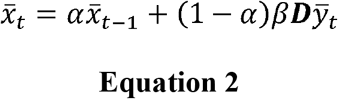

In Eq. 2, 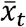 denotes a 2*F*_*N*_ ×1 column vector of position and velocity estimates for *F*_*N*_ fingers, ***D*** is a *E*_*N*_ × 2*F*_*N*_ matrix for *E*_*N*_ electrodes, α is the scaling factor, and β is the gain factor. For the position-velocity Kalman filter, the smoothing factor and gain were only implemented for the velocity kinematics. To determine the optimal values for α and β, one day of testing with RK was dedicated to first tuning the gain, β, by increasing its value. Using the value of β giving the best performance, the smoothing factor, α, was then adjusted. The optimal values, based on throughput, were determined to be α = 1.25 and β = 1.2. The performance improvements from using a zero-lag RK and optimally tuned gain and smoothing factors were tested on a second day of testing. Using RK with optimally tuned parameters (475 trials) resulted in only a 3.6% increase in throughput and 16% increase in average peak finger velocity.

Although most comparisons of RN and RK use RK without tuned hyperparameters, we did include a test RN and RK_opt_, which uses zero lag and optimal values of α = 1.25 and β = 1.2, and validated our findings with a fully optimized RK at 29 mos post implantation (see Results Section “ReFIT neural network decoder outperforms both the original NN and RK decoders”).

## Acknowledgments

We thank Eric Kennedy and Paras Patel for animal and experimental support. We thank Gail Rising, Amber Yanovich, Lisa Burlingame, Patrick Lester, Veronica Dunivant, Laura Durham, Taryn Hetrick, Helen Noack, Deanna Renner, Michael Bradley, Goldia Chan, Kelsey Cornelius, Courtney Hunter, Lauren Krueger, Russell Nichols, Brooke Pallas, Catherine Si, Anna Skorupski, Jessica Xu, and Jibing Yang for expert surgical assistance and veterinary care. We thank Tom Cichonski for his editorial review.

This work was primarily supported by NSF grant 1926576. M.S.W., H.T., M.J.M. were supported by NSF grant 1926576. S.R.N. was supported by NIH grant F31HD098804. J.T.C. was supported by NSF GRFP. P.G.P. was supported by NSF grant 1926576, the A. Alfred Taubman Medical Research Institute, and NIH grant R01GM111293. C.A.C. was supported by NSF grant 1926576, Craig H. Neilsen Foundation project 315108, NIH grant R01GM111293, and MCubed project 1482. This work was also supported in part by NIH grant R01GM111293.

## Author contributions

M.S.W., S.R.E., H.T., M.J.M., and J.T.C. performed the NHP decoding experiments. M.S.W. conducted the data analysis and wrote the manuscript. M.S.W. and S.R.N. wrote and validated the analysis software. M.S.W. and S.R.E. wrote and validated the Python code. S.R.E. developed and tested the xGPC. M.S.W., P.G.P., and C.A.C. conducted the surgeries. P.G.P. and C.A.C. supervised this work. All authors reviewed and modified the manuscript.

## Videos

**Video 1** | **RK for a random finger task**. The video captures the average performance of the RK as measured by throughput (1.4 bps) over the two days of testing for Monkey N when comparing RK and RN.

**Video 2** | **RN for a random finger task**. The video captures the average performance of the RN decoder as measured by throughput (2.3 bps) over the two days of testing for Monkey N when comparing RK and RN.

**Video 3** | **RN for a center-out finger task**. The video captures the peak performance of the RN decoder as measured by throughput (3.2 bps) in one day of testing of the RN for Monkey N on a typical center-out finger task.

